# Deterministic co-encapsulation of microparticles in droplets via synchronized merging for single-cell genomics

**DOI:** 10.1101/2025.10.10.681553

**Authors:** Yanan Du, Chuhe Wen, Yuan Luo, Yifan Liu

## Abstract

Droplet encapsulation of microparticles such as hydrogel beads and cells is fundamental to high-throughput bioanalytical assays, including single-cell genomics. However, co-encapsulation of multiple particles at defined ratios remains a technical challenge. Here, we present a simple and robust strategy for deterministic co-encapsulation, combining particle-triggered droplet generation with synchronized droplet merging. Close-packed hydrogel particles initiate droplet formation, and the resulting droplets are synchronized and merged within a microfluidic channel, enabling efficient encapsulation of microparticles at user-defined ratios. We systematically characterized the process and demonstrated reliable co-loading of 55 µm and 80 µm hydrogel particles at 1:1 and 2:1 ratios. Using this platform, we developed a single-cell 16S rRNA gene sequencing workflow by co-encapsulating hydrogel capsules containing amplified microbial genomes with barcode beads. Applied to a 1:1 mixture of *Escherichia coli* and *Bacillus subtilis*, the method enabled accurate, high-throughput single-cell taxonomic profiling. This co-encapsulation approach offers broad applicability for single-cell genomics, digital assays, and interaction-based studies requiring precise particle pairing.

## INTRODUCTION

Droplet microfluidics enables the high-throughput encapsulation and processing of micrometer-scale particles within picoliter to nanoliter partitions^1^. This approach has driven significant innovations in bioanalytical chemistry, including single-cell genomics^2, 3^, digital immunoassays^4^, and high-throughput screening^5, 6^. Typically, single microparticles (e.g., cells or beads) are loaded into droplets via limiting dilution, a process governed by Poisson statistics and yielding an encapsulation efficiency of approximately 5%^7^. In many applications, this low efficiency is offset by the extremely high droplet generation rates^3, 8^. To improve encapsulation efficiency, microfluidic strategies such as inertial ordering^9^ and hydrogel close-packing^10^ have also been widely employed.

As droplet-based biotechnologies continue to evolve, many applications require the encapsulation of more than one particle in a droplet^1^. For instance, in single-cell genomics, a barcode bead must be paired with either a cell^10^ or a processed genome housed in a gel bead^11^. Some methods demand the encapsulation of more than two particles to facilitate interaction-based analyses. For example, Gérard et al.^12^ reported the CelliGO system, a microfluidics-based platform designed to characterize the antibody phenotype and sequence at the single-cell level by co-encapsulating immune cells, cancer cells, and barcode beads. However, the efficiency of multi-particle loading is often limited by the overlapping effects of multiple Poisson distributions, resulting in low co-encapsulation rates^12^. This limitation highlights the need for more effective strategies. Current deterministic approaches often rely on active droplet manipulation, such as fluorescence-based sorting^13^, which adds complexity and reduces throughput. An alternative, passive strategy exploits the “zipping” behavior of two converging streams of close-packed hydrogel beads^14^; however, this method imposes strict constraints on bead size and achievable co-encapsulation ratios. Consequently, achieving rapid and reliable co-loading of multiple microparticles—particularly those with differing sizes or material properties—remains a significant technical challenge.

Here, we present a microfluidic strategy for high-efficiency co-encapsulation of particles. The method comprises two sequential steps: (1) particle-triggered droplet generation and (2) droplet merging. As illustrated in Fig. 1a, the first step produces particle-containing droplets with >90% efficiency. This is achieved by introducing close-packed elastic hydrogel particles into a dripping aqueous jet. By carefully tuning the flow rates, the particle loading frequency matches the rate of droplet formation. Notably, the presence of a particle within the jet increases the local curvature, promoting jet breakup via surface tension– driven Rayleigh-Plateau instability and further enhancing encapsulation efficiency. The resulting droplets are slightly larger than the encapsulated particles. In the second step (Fig. 1b), droplets are paired at defined ratios and merged, enabling deterministic co-encapsulation of multiple particles. Both steps are integrated into a single microfluidic device, and each was thoroughly characterized. Using this approach, we developed a single-cell 16S sequencing method that co-encapsulates a hydrogel capsule^15, 16^ containing an amplified microbial genome with a barcode bead, followed by indexing PCR within droplets. We demonstrate that this method enables high-throughput, single-cell profiling of 16S rRNA gene sequences from a mixed population of *E. coli* and *B. subtilis*. This co-encapsulation strategy offers substantial potential for scalable biotechnological applications, including single-cell analysis and digital assays.

**Figure 1.**
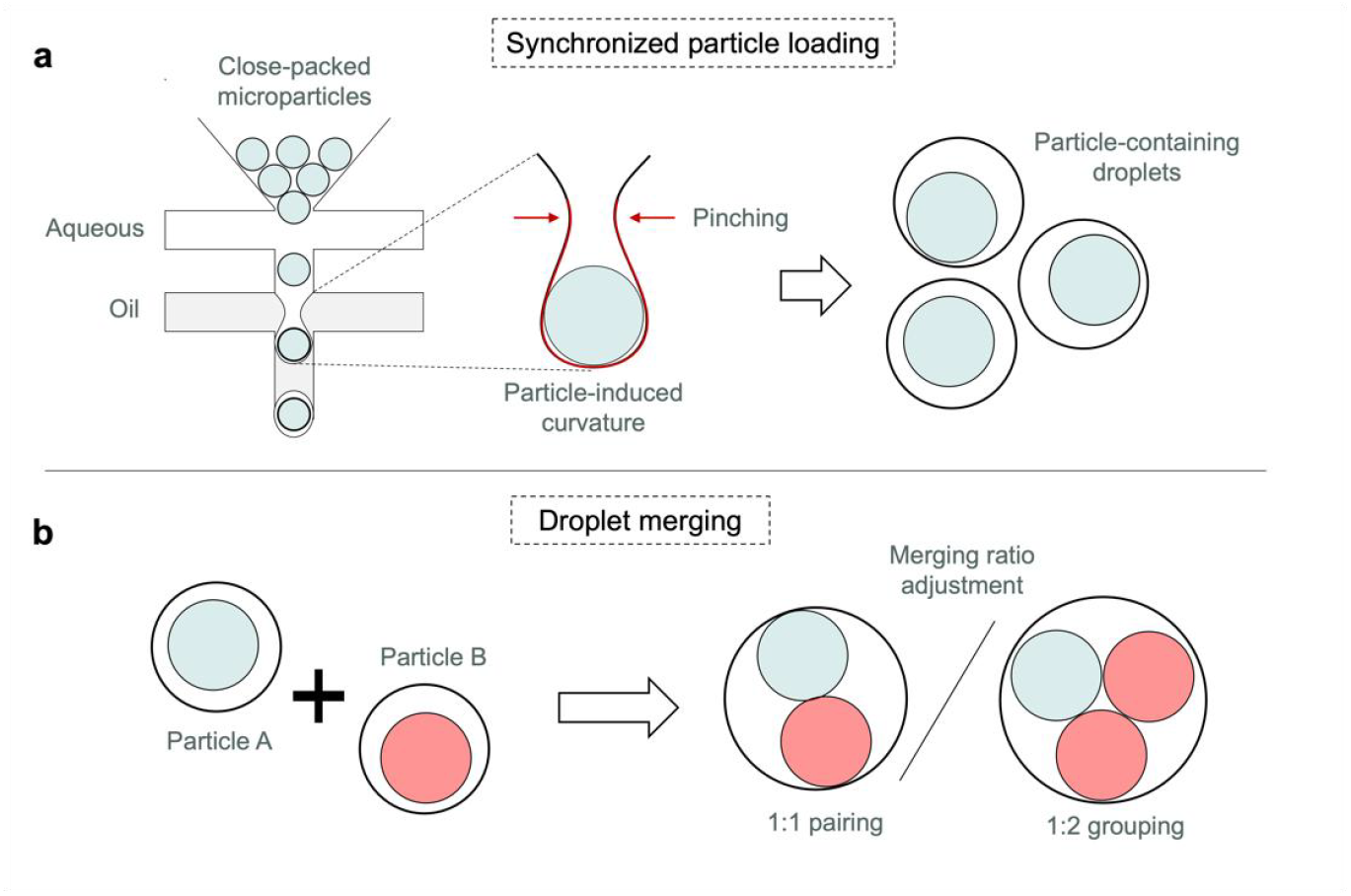
Conceptual illustration of the deterministic encapsulation workflow. (a) Hydrogel microparticles are synchronously loaded into droplets, resulting in particle-containing droplets with high efficiency. (b) Droplets containing different microparticles are paired and merged, enabling co-encapsulation at defined ratios.

## RESULTS AND DISCUSSIONS

### Synchronized loading of hydrogel particles

Close packing of hydrogel beads was initially reported by Abate et al. as a means to improve encapsulation efficiency^17^. A more recent work has demonstrated particle-triggered droplet generation in the jetting regime, further enhancing the encapsulation performance^18^. In this study, we sought to leverage these mechanisms within the dripping regime, where droplet formation is more periodic and controllable—an advantage for downstream processes such as droplet pairing and merging. To characterize the process, we approximated a two-dimensional droplet generation geometry, as illustrated in Fig. 2a. This configuration consisted of two adjacent microfluidic junctions: the first used a pinching aqueous flow to space close-packed hydrogel beads at regular intervals, while the second introduced an oil phase to generate droplets.

**Figure 2.**
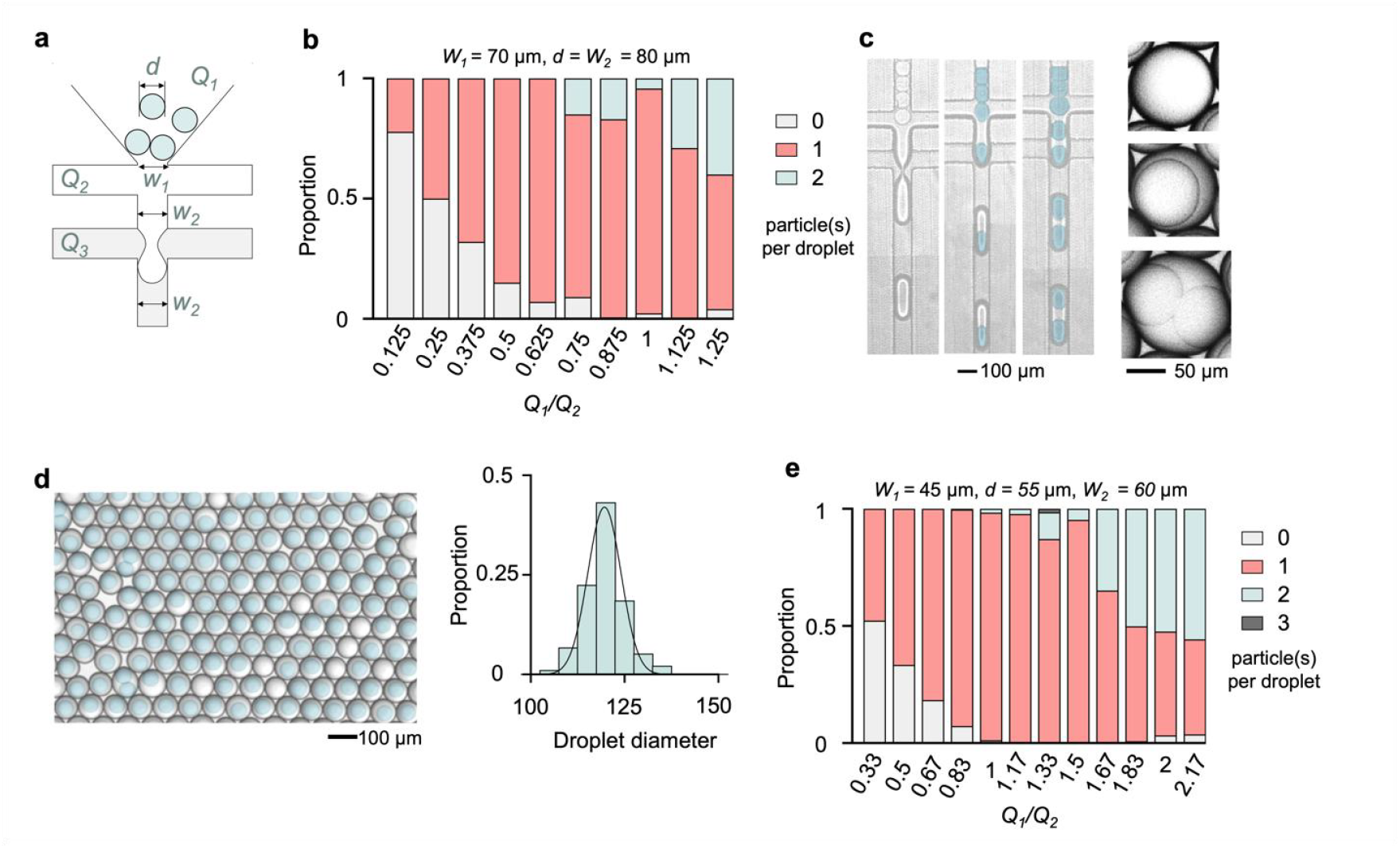
Characterization of the particle loading process. (a) Schematic of a dual-flow-focusing junction for particle loading. Key parameters: bead-loading flow rate (*Q*_*1*_), aqueous spacer flow rate (*Q*_*2*_), oil flow rate (*Q*_*3*_), width of the first junction (*W*_*1*_), width of the second junction (*W*_*2*_), particle diameter (*d*). (b) Particle (d = 80 µm) loading behavior at varying *Q*_*1*_/*Q*_*2*_ values (n > 300). (c) Micrographs depicting the loading process and particle-containing droplets for the particle loading experiments in (b). (d) Droplets and their size distributions (n = 300) of a representative particle loading experiment (*Q*_*1*_/*Q*_*2*_ = 1). (e) The loading behavior for 55 µm particles at varying *Q*_*1*_/*Q*_*2*_ values (n > 300).

A critical prerequisite of bead close packing is that the entrance width (*W*_*1*_) must be slightly smaller than the bead diameter (*d*). Moreover, the width of the second junction (*W*_*2*_) should be comparable to *d* to ensure an appropriate size of the resulting droplets. Considering these, we first tested with *W*_*1*_ of 70 μm and *W*_*2*_ of 80 μm (Fig. S1a), and evaluated the loading of 80 μm polyacrylamide hydrogel beads (Fig. 2b). In the experiments, we fixed the aqueous and oil flow rates (*Q*_*2*_ = 400 µL/h and *Q*_*3*_ = 800 µL/h), and monitored the ratio between bead injection flow and aqueous spacer flow (*Q*_*1*_*/Q*_*2*_). The result suggested a clear dependence of bead loading efficacy on *Q*_*1*_*/Q*_*2*_. The best loading performance was found at *Q*_*1*_*/Q*_*2*_ = 1, where 93.5% of the resulting droplets contained one bead. Such a strong and quasi-linear dependence (Fig. S2a) of bead loading on *Q*_*1*_*/Q*_*2*_ (more specifically *Q*_*1*_, as *Q*_*2*_ is fixed) indicated that although the particle flow varied, the resulting droplet size was more or less maintained.

We then looked at three specific *Q*_*1*_*/Q*_*2*_ values (0.125, 0.625, 1.25) where the bead loading behavior was distinct (Fig. 2c). At *Q*_*1*_*/Q*_*2*_ = 0.125 where most droplets were empty, the average droplet size was 109 µm (Fig. S2b). At *Q*_*1*_*/Q*_*2*_ = 0.625 where ~90% contained a bead, the size was 120 µm (Fig. S2c). This observed tolerance of droplet size on the discrete flow closely agrees with the droplet dripping characteristics. In both scenarios, the bead-containing droplets were 5% larger than the empty ones (Figs. S2b&c). This suggests that bead loading indeed influenced the droplet dripping dynamics, as we proposed in Fig. 1a.

Based on the above results, we selected the condition of *Q*_*1*_*/Q*_*2*_ = 1 for further characterization. We looked into the monodispersity of the resulting droplets (Fig. 2d) and found that the droplet sizes were tightly distributed around 118 μm (S.D. = 3.79). To test whether *Q*_*1*_*/Q*_*2*_ = 1 is a universal parameter for efficient bead loading, we fabricated another loading geometry with *W*_*1*_ = 45 μm and *W*_*2*_ = 60 μm and loaded 55 μm beads (Fig. S1b). As shown in Fig. 2e, we again obtained the best loading efficiency at *Q*_*1*_*/Q*_*2*_ = 1, where 97% of the droplets encapsulated one bead.

### Co-encapsulation of hydrogel particles

Following optimization of the loading process, we designed and fabricated a microfluidic device to enable pairing and merging of particle-loaded droplets. The device consists of two particle-loading modules and a downstream droplet merging region (Fig. 3a and Fig. S1c). Droplets generated at matched frequencies from the two modules are paired as they flow downstream and subsequently merged at a serrated “teeth” structure under an applied electric field. Pairing is achieved through a size asymmetry between the droplet populations. In our design, droplets containing particle A are smaller than those containing particle B. As a result, “A” droplets remain narrower than the channel width, while “B” droplets span the full width. Under pressure-driven flow, the smaller A droplets are centered within the channel by shear-induced lift forces^19^ and travel faster than the larger B droplets, leading to effective droplet pairing. At the merging site (Fig. 3a, inset micrograph), coalescence is driven by a combination of dielectrophoresis, electrostatic interactions, capillary instabilities, and interface destabilization^20^. Specifically, the applied AC field induces temporary dipoles within the droplets, generating electrostatic attraction and dielectrophoretic forces that align and draw droplets together. Concurrently, the field deforms the droplet interfaces, thinning the intervening film. Electrohydrodynamic flows and interfacial instabilities further disrupt this film, ultimately leading to its rupture and droplet merging (Fig. 3a inset).

**Figure 3.**
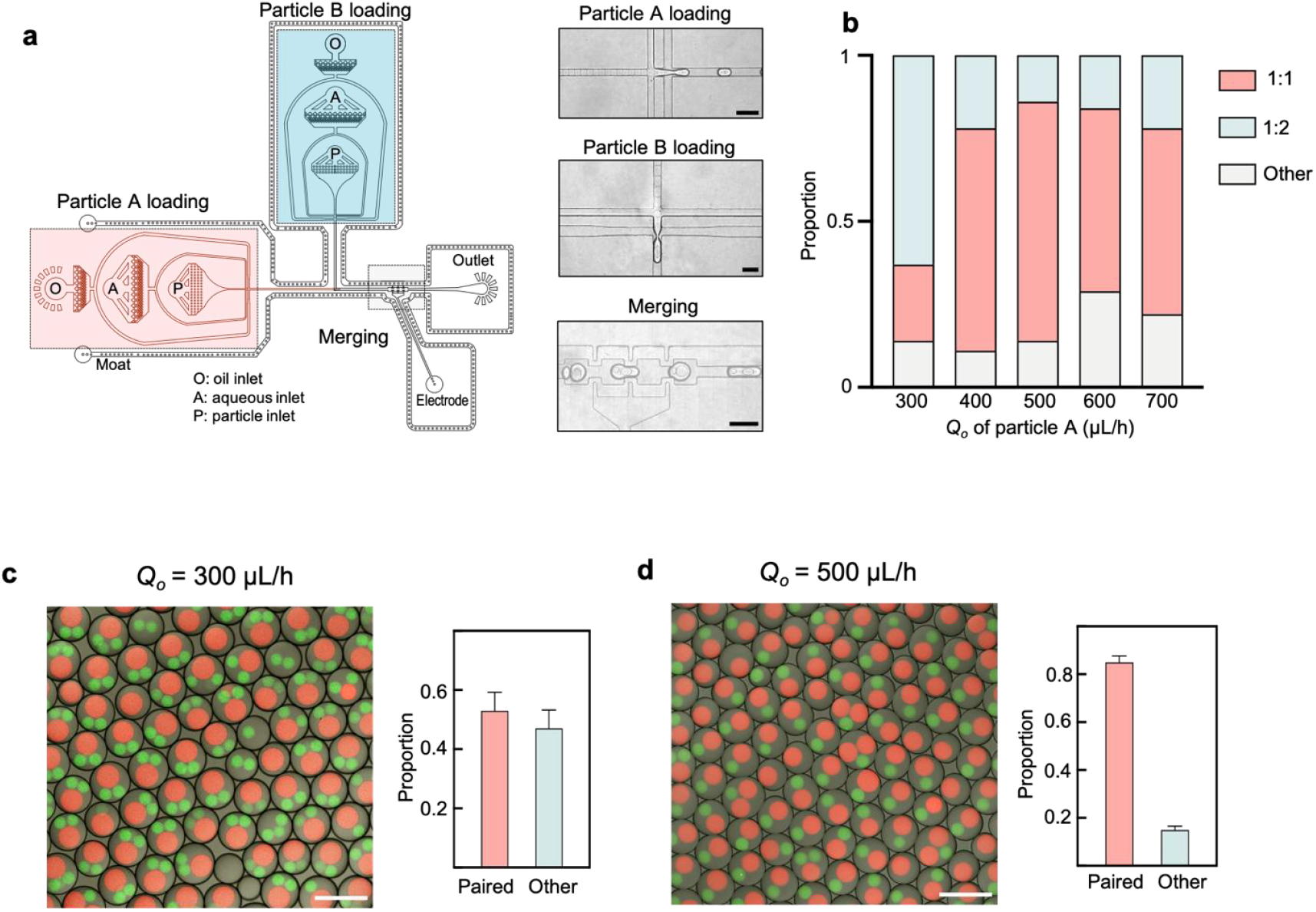
Characterization of the particle pairing and merging. (a) Layout of a microfluidic design consisting of two particle loading modules and a droplet merging geometry (see insets for micrographs) (Scale bar: 200 µm). (b) f. (*Q*_*o*_). (c&d) Representative co-encapsulation experiments and statistics conducted at (c) *Q*_*o*_ = 300 µL/h (n = 552) and (d) *Q*_*o*_ = 300 µL/h (n = 816) (Scale bar: 200 µm).

To characterize the co-encapsulation process, we used hydrogel particles of 55 μm (particle A) and 80 μm (particle B). The flow parameters for particle B were fixed (total aqueous flow: 650 µL/h; oil: 800 µL/h), while the oil flow rate for particle A (*Qo*) was varied (Fig. 3b). At a low oil flow rate (*Qo* = 300 µL/h), droplets predominantly formed 2:1 groupings (62.6%; Fig. 3c). Analysis of droplet movement dynamics (Supplementary Movie 1) revealed that particle A droplets were spaced more closely than particle B droplets—approximately half the inter-droplet distance. We next increased the oil flow rate for particle A and found it reduced the frequency of 2:1 group formation but concurrently increased the occurrence of unpaired droplets (0:1). Reasonable 1:1 pairing was achieved at *Qo* = 500 µL/h, where 72.2% of droplets were successfully paired in a 1:1 ratio (Fig. 3d). Figs. 3c and 3d show representative micrographs of co-encapsulated particles (green: particle A; red: particle B) at two distinct *Qo* values, along with corresponding statistical analyses. To further optimize pairing, we varied the oil flow rate for particle B between 700 and 1100 µL/h, achieving a maximum 1:1 pairing efficiency of 90.6% (Fig. S3a). Notably, paring two particle B and one particle A is also feasible (Fig. S3b). These results demonstrate the feasibility of deterministic co-encapsulation through precise control of flow conditions.

### Single-cell 16S rRNA gene sequencing

Having optimized the co-encapsulation strategy, we developed a single-cell 16S rRNA gene sequencing workflow based on this approach. The workflow comprises two main steps. In the first step, individual microbial cells are encapsulated in hydrogel microcapsules (70 um in diameter), followed by cell lysis and whole-genome amplification. This step was adapted from CAP-seq^15^, a method we previously developed for sequencing the whole genomes of individual microbial cells. In the second step, the capsules containing amplified genomes are co-encapsulated with barcode beads in droplets (Fig. 4a). This is achieved using the same device in Fig. 3a with optimized fluidic parameters (see Methods). A barcoding PCR is then performed within the droplets to amplify and index the V1–V3 region (507 bp) of the 16S rRNA genes. Finally, the barcoded PCR products (578 bp) were purified, subjected to library preparation, and sequenced on a nanopore sequencer.

**Figure 4.**
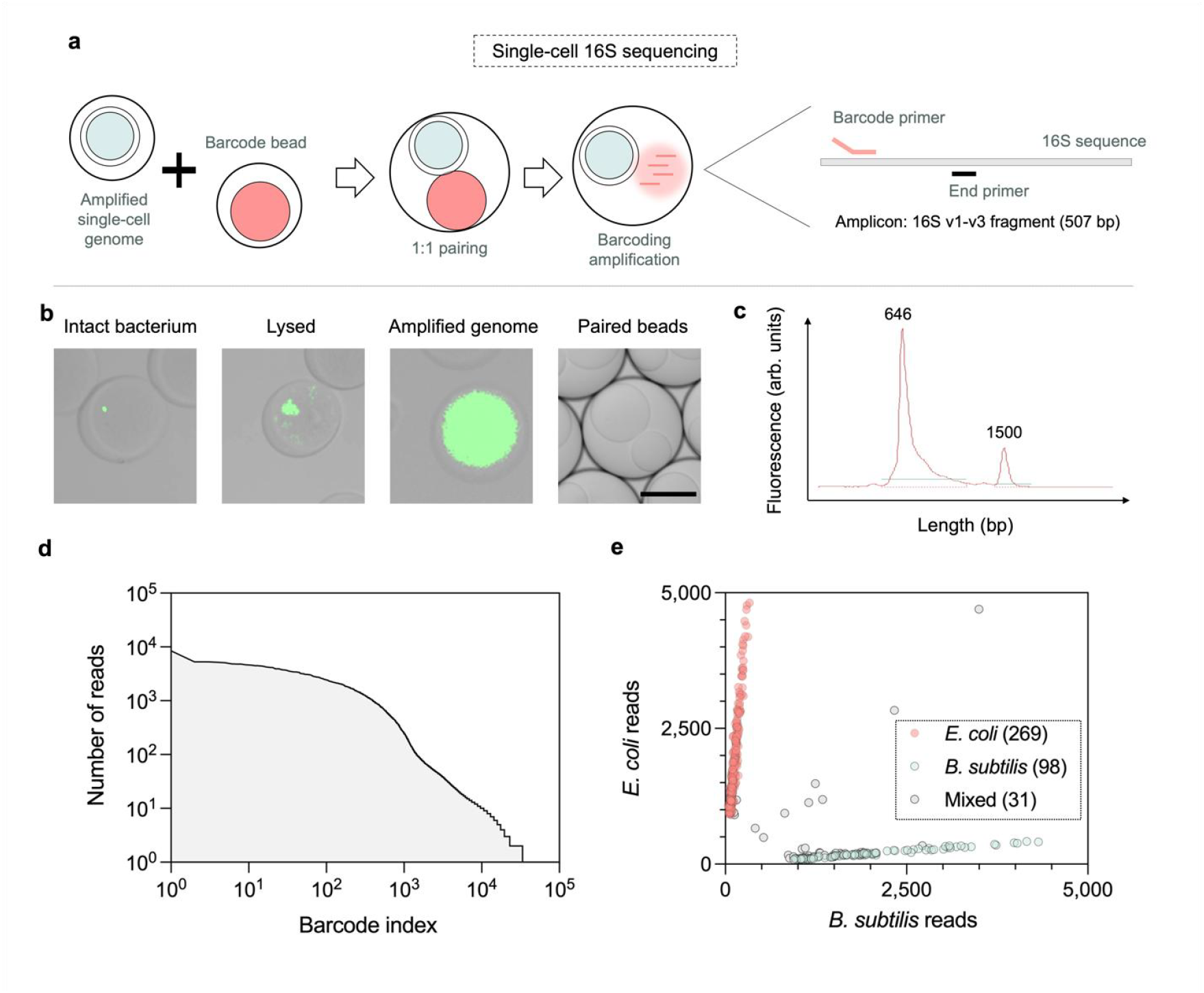
Developing single-cell 16S rRNA gene sequencing with the deterministic co-encapsulation strategy. (a) Conceptual illustration of the sequencing workflow. Hydrogel capsules containing amplified single-cell genomes are co-encapsulated with barcode beads to selectively index and amplify v1-v3 region of the 16S rRNA gene. (b-e) Sequencing a mixture of *E. coli* and *B. subtilis*: (b) Micrographs depicting the key steps of the sequencing workflow. Scale bar: 50 µm. (c) DNA length profile of the barcoded amplicon library (theoretical length: 578 bp). The 1500 bp peak denotes a DNA marker. The result was generated by an Agilent Bioanalyzer using a DNA 1000 kit. (d) Barcode group profile ordered by the number of reads of each barcode group. (d) Scatter plot depicting 398 barcode groups whose reads are mapped to the two references.

As shown in Fig. 4b, we validated the key workflow steps—including cell encapsulation, cell lysis, whole-genome amplification, and pairing of capsules with barcode beads—using fluorescence microscopy and DNA staining. The capsule–barcode bead pairing achieved an efficiency of approximately 80%. We then analyzed the size distribution of the barcoded PCR products (Fig. 4c), which exhibited a single peak at ~650 bp, closely matching the expected product size of 578 bp.

Next, we evaluated the single-cell 16S rRNA sequencing workflow using a 1:1 mixture of *E. coli* and *B. subtilis* cells. The sequencing run generated 2,676,425 reads with an average read length of 753 bp (Fig. S4). Subsequent bioinformatic analysis identified 398 unique barcode groups containing more than 1,000 reads each (Fig. 4d). To assess whether each barcode group represented an individual cell, we mapped the reads within each group to reference 16S rRNA gene sequences of *E. coli* and *B. subtilis*. A barcode group was assigned to a species only if more than 95% of its reads mapped to that species; otherwise, it was categorized as “mixed.” As shown in Fig. 4e, 269 barcode groups were assigned to *E. coli* and 98 to *B. subtilis*. The observed doublet rate was 7.79%. These results clearly demonstrate that the workflow enables 16S rRNA gene analysis with single-cell resolution.

In conclusion, we have demonstrated a robust method for co-encapsulating microparticles in droplets at defined ratios, using a single microfluidic device that integrates two sequential steps: (1) particle-templated droplet generation and (2) droplet merging. While we showcased 1:1 and 1:2 pairings of two distinct microparticles, the method is adaptable to a wider range of pairing ratios and additional particle types through flow rate adjustments or incorporation of extra particle-loading modules^5^. To further demonstrate the novelty and capabilities of the deterministic co-encapsulation of microparticles technology, we have provided a multi-dimensional comparative analysis between our method and several representative bead-loading approaches, including In-Drop^10^, CelliGo^12^, and the microfluidic “zipper”^14^ (Table S1). Beyond hydrogel particles, the strategy is compatible with other materials—such as rigid microbeads and cells—via a hydrogel-coating approach^7^. Therefore, this approach should facilitate interaction-based analyses involving multiple microparticle types, such as sequencing paired T cells and cancer cells^12, 13^. To desmonstrate its broader utility, we applied the platform to bacterial systems by encapsulating bacteria expressing green or red fluorescent proteins into distinct hydrogel beads. Co-encapsulation of these beads within single droplets created a controlled microenvironment for investigating bacterial interactions, including interspecies communication, metabolic cooperation, and antagonism behaviors (Fig. S5).

In the present study, we reported a high-throughput single-cell 16S rRNA gene sequencing technique enabled by facilitating the precise pairing of hydrogel capsules containing amplified microbial genomes with barcode beads. The 16S rRNA gene serves as a molecular fingerprint for bacterial identification, providing species- and sometimes strain-level resolution based on its conserved and variable regions. The developed single-cell approach therefore enables high-resolution analysis of microbial diversity and heterogeneity by capturing genetic information from individual cells without reliance on population-level averages. Beyond 16S rRNA genes, the method is readily adaptable to other gene targets or gene sets through modification of primer designs. For example, it can be used to detect antibiotic resistance genes at the single-cell level, revealing the distribution of resistance determinants across individual bacteria within complex communities.

While the platform demonstrates high encapsulation fidelity and versatility, several current limitations and future directions warrant consideration. For instance, scaling up the platform for larger-scale applications may encounter challenges such as the limited number of hydrogel beads that can be reliably loaded into syringe, as well as potential microchannel clogging under high particle concentrations. In addition, the reliance on microfluidic devices and specialized operational skills increases the technical difficulty, limiting widespread option. Future work will therefore focus on optimizing chip architecture, incorporating active flow control, and exploring parallelization strategies to enhance throughput and robustness. Taken together, we anticipate that the reported approach will be broadly applicable to the field of single-cell genomics.

## MATERIALS AND METHODS

### Device fabrication

The microfluidic device was fabricated using PDMS-based soft lithography. The layout was designed in AutoCAD and printed as dark-field plastic film masks. A negative photoresist (SU-8 3025, MicroChem) was spin-coated onto a 3-inch silicon wafer, exposed to UV light, and developed following the manufacturer’s instructions. The mold was cleaned with isopropanol and ethanol, then blow-dried using nitrogen gas. The PDMS precursor (SYLGARD 184, Dow Corning) was mixed with its curing agent (10:1, w/w) and degassed. After pouring the PDMS over the mold, it was cured at 60 °C overnight. The cured PDMS was peeled from the mold, and inlet/outlet ports were created using a 0.7-mm hole puncher. The slab was bonded to a clean glass slide using oxygen plasma treatment and baked at 100 °C for 30 minutes. The channels were then made hydrophobic by treating the device with Aquapel (PPG Industries). A high-resolution layout is provided in the Supporting Information.

### Hydrogel bead production

The hydrogel precursor mixture consisted of 6% (w/v) acrylamide, 0.15% (w/v) N,N’-bis(acryloyl)cystamine, 0.3% (w/v) ammonium persulfate (APS), and 1 µM acrydite-modified oligonucleotides (particle A: CCAAGCAGAAGACGGC–Acrydite; particle B: GCTTACGAGACCGGA–Acrydite). This hydrogel solution was used as the dispersed phase, while the continuous phase comprised HFE-7500 containing 2% (w/v) PEG-PFPE surfactant. Hydrogel particle A (55 µm) was generated using a standard flow-focusing device with a channel width of 40 µm and height of 27 µm, with flow rates of 450 µL/h for the continuous phase and 200 µL/h for the dispersed phase. Particle B (80 µm) was produced using a chip with the same channel width (40 µm) but a height of 50 µm, with corresponding flow rates of 600 µL/h (continuous phase) and 300 µL/h (dispersed phase). To initiate gel polymerization, 1% (v/v) N,N,N’,N’-tetramethylethylenediamine (TEMED) was added to the droplets, followed by incubation at room temperature for 4 hours. Hydrogel particles were recovered by adding 20% (v/v) 1H,1H,2H,2H-perfluoro-1-octanol in HFE-7500, vortexing, centrifugation, and removal of the oil phase by pipetting. The particles were washed three times in 0.1% Tween-20 in Dulbecco’s phosphate-buffered saline (DPBS), which also served as the storage buffer. Subsequently, fluorescently labeled microspheres were prepared via a hybridization reaction. The hybridization master mix contained 1 µM probe oligonucleotides (particle A: GCCGTCTTCTGCTTGG–FAM; particle B: TCCGGTCTCGTAAGC–Cy3), 5 µg/mL BSA, and 0.4 × phosphate-buffered saline (PBS). Thermal cycling was carried out using the following conditions: 68 °C for 5 min, 48 °C for 8 min, 40 °C for 8 min, 30 °C for 8 min, followed by a hold at 4 °C. The particles were then washed three times in TET buffer (10 mM Tris-HCl, pH 8.0; 10 mM EDTA; 0.1% Tween-20), which also served as the final storage buffer.

### Microfluidic experiments

The prepared hydrogel beads were first pelleted by centrifugation at 4000 rpm for 1 minute to remove excess buffer. The concentrated beads were then transferred to a 1 mL syringe for reinjection. Hydrogel beads A were reinjected into the left inlet of a dual-junction flow-focusing device (Fig. S1a.) at varying flow rates ranging from 40 to 260 µL/h. The aqueous flow rate was fixed at 120 µL/h, while the carrier oil flow rate was varied between 300 and 700 µL/h. Similarly, hydrogel beads B were reinjected into the top inlet of a dual-junction flow-focusing device (Fig. S1b.) at flow rates ranging from 50 to 500 µL/h. The aqueous flow rate was also fixed at 120 µL/h, and the carrier oil flow rate was varied between 300 and 700 µL/h. The aqueous phase in all reinjection experiments consisted of 1× phosphate-buffered saline (PBS). For particle pairing, the flow rate of hydrogel beads A and B fixed at 100 µL/h and 250 µL/h respectively, while the responding aqueous flow rate was fixed at 120 µL/h (A) and 400 µLh (B). Firstly, the carrier oil flow rate for hydrogel beads B was fixed at 800 µL/h, while the carrier oil (for beads A) flow rate was varied between 300 and 700 µL/h. Then, the carrier oil flow rate for hydrogel beads A was fixed at 500 µL/h, while the carrier oil (for beads B) flow rate was varied between 700 and 1100 µL/h. At the micro-teeth structure, the grouped droplets were merged under an AC electrical field (1.5 kV Vpp, 50 kHz) generated by an inverter (TDK CXA-L0605-VJL) circuit. The moat region surrounding the merging site was filled with 2 M KCl solution, which provides sufficient conductivity to ensure efficient field generation while maintaining the integrity of droplets. For single-cell 16S sequencing, barcode beads were reinjected into the left inlet of a dual-junction flow-focusing device at flow rates of 120 µL/h. The aqueous (nuclease-free water) flow rate was fixed at 120 µL/h, while the carrier oil flow rate was 500 µL/h. Similarly, CAPs were reinjected into the top inlet of a dual-junction flow-focusing device at flow rates ranging of 250 µL/h. The aqueous (PCR mix: 1.7 × Phusion High-Fidelity reaction buffer (NEB, M0530L), 1.7 mM dNTP, 17 mM DTT, and 0.85 U/µL Phusion High-Fidelity DNA Polymerase (NEB, M0530L)) flow rate was fixed at 400 µL/h, and the carrier oil flow rate was 1000 µL/h.

### Cell culture

*Escherichia coli* (strain MG1655), *Bacillus subtilis* (strain 168), and *Escherichia coli* (strain BL21) expressing green or red fluorescent proteins were cultured overnight at 37°C in Luria-Bertani (LB) medium. Then, 1 mL of the bacterial cultures was washed three times in 1 mL DPBS before resuspending in 1 mL DPBS. Cell concentration is determined by manually counting serial dilutions of the liquid culture on plastic slides (Thermo Fisher, C10228) using a microscope. The *Escherichia coli* (strain MG1655) and *Bacillus subtilis* (strain 168) cell suspension was then used as input for single-cell 16S sequencing. While the cell suspension of *Escherichia coli* (strain BL21) expressing green or red fluorescent proteins cell suspension were separately encapsulated into distinct hydrogel beads and subsequently co-encapsulated within single droplets.

### Single-cell 16S rRNA gene sequencing

Single-cell 16S sequencing was performed using a workflow adapted from CAP-seq, a previously developed method for whole-genome sequencing of single microbes. A 1:1 mixture of *E. coli* and *B. subtilis* cells was encapsulated in hydrogel capsules using a microfluidic device. Cell lysis was carried out in two steps: (1) incubation in a buffer containing 10 mM Tris-EDTA, 50 mM NaCl, 5 mM EDTA, 0.5% Triton X-100, 900 U lysozyme, and 10 U lysostaphin at 37 °C for 2 hours; followed by (2) incubation in 50 mM NaCl, 5 mM EDTA, 0.5% SDS, and 0.5 mg/mL proteinase K. The resulting capsules, containing exposed genomes, were incubated in a multiple displacement amplification (MDA) mix for whole-genome amplification. Capsules with amplified genomes were then co-encapsulated with barcode beads (CTACACGACGCTCTTCCGATCTNNNNNNNACAGNNNNNNNGTCANNNNNNNAGAGTTTGA TCMTGGCTCAG) and an end primer (ATTACCGCGGCTGCTGG) to amplify the V1–V3 region of the 16S rRNA gene. The final buffer conditions in the droplets were 1 × Phusion High-Fidelity reaction buffer (NEB, M0530L), 1 mM dNTP, 10 mM DTT, and 0.05 U/µL Phusion High-Fidelity DNA Polymerase (NEB, M0530L). The thermal cycling was performed using the following parameters: 98°C for 3 min, 15 cycles of 98 °C for 10 s, 55 °C for 30s and 72°C for 30 s, 72°C for 5 min, and 4 °C hold. Amplified DNA products were extracted and purified as previously described^15^. The library was characterized on an Agilent Bioanalyzer using a DNA 1000 kit (Fig. 4c). Sequencing libraries were prepared according to the manufacturer’s instructions and sequenced on a nanopore sequencer (Oxford Nanopore PromethION 2).

### Hydrogel bead containing bacteria preparation

The *Escherichia coli* (strain BL21) expressing green or red fluorescent proteins cell is resuspended by the 1 mL hydrogel precursor mixture consisted of 6% (w/v) acrylamide, 0.15% (w/v) N,N’-bis(acryloyl)cystamine, and 0.3% (w/v) ammonium persulfate (APS) to a final concentration of ~5 cells/bead. Then the hydrogel beads containing cells is collected according to the previous method.

### Bioinformatics

The raw sequencing data were deposited on the NCBI database with an accession number of PRJNA1227260. The sequencing reads were first aligned to a known adapter sequence (AGAGTTTGATCATGGCTCAG) using the Smith-Waterman algorithm, with barcodes identified based on alignment scores of 30 or higher. Verified barcodes were then added to the read names for subsequent demultiplexing. Next, the adapter and surrounding barcode sequences were removed using the Cutadapt tool, leaving clean 16S sequences. These reads were demultiplexed using BBMap’s demuxbyname.sh script, which generated separate FASTQ files for each barcode. A knee plot of the cumulative read count distribution was generated, and only barcodes with more than 1,000 reads were kept. The remaining reads were aligned to the 16S V1-V3 region sequences of *E. coli* MG1655 (with the *rrsH* gene as reference) and *B. subtilis* strain 168 using Minimap2 with the parameters -ax map-ont - secondary=no (default settings for other parameters).

## Notes

### Competing Interest Statement

The authors have declared no competing interest.

