## Supplementary materials for "Deterministic co-encapsulation of microparticles in droplets via synchronized merging for single-cell genomics"

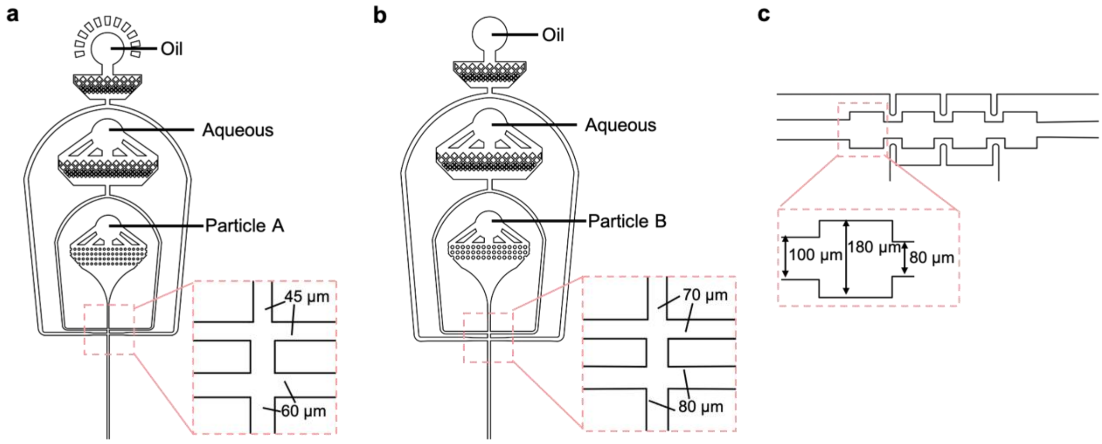


**Figure S1.** The layout of microfluidic devices/structures for (a) generating particle A-encapsulated droplets and (b) particle B-encapsulated droplets and (c) droplet merging.

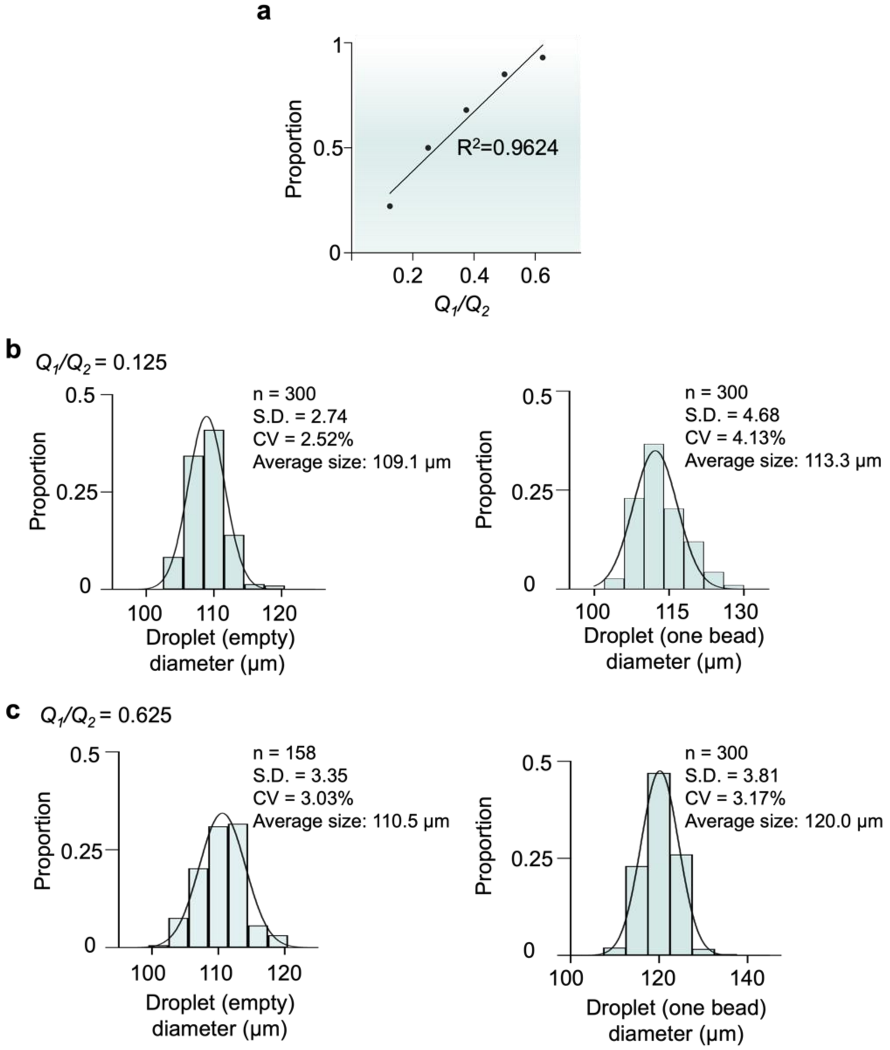


**Figure S2.** (a) A quasi-linear relationship observed between the proportion of droplets containing one bead and the ratio between bead injection flow and aqueous spacer flow ($Q_1/Q_2$). The solid line depicts a linear fit of the scattered dots. (b&c) Size distribution of droplets (left: empty droplets; right: particle-containing droplets) generated at (b) $Q_1/Q_2$ = 0.125 and (c) $Q_1/Q_2$ = 0.625. The curves exhibit the Gaussian fitting results.

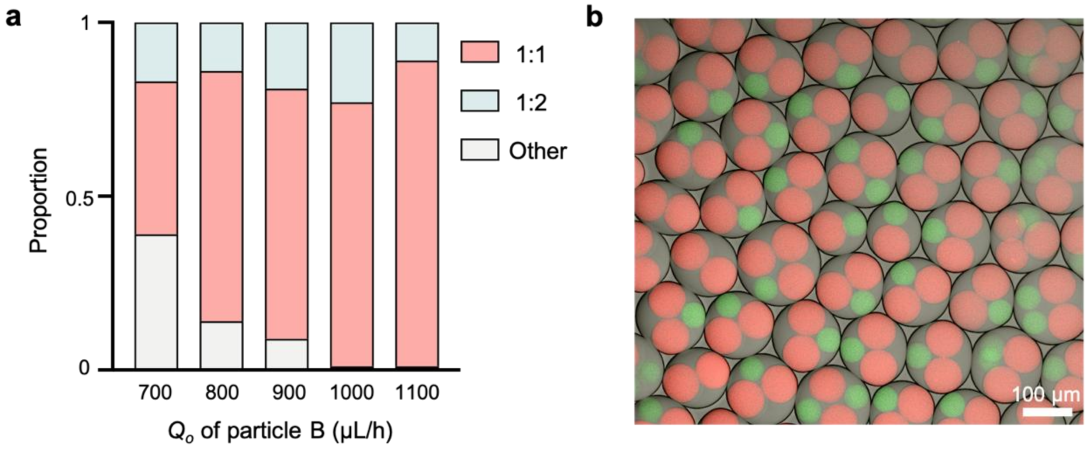


**Figure S3.** (a) Statistics of co-encapsulation at different oil flow rates ($Q_o$) for particle B. (b) Representative co-encapsulation of two paticle A (80 µm) beads with one particle B (55 µm).

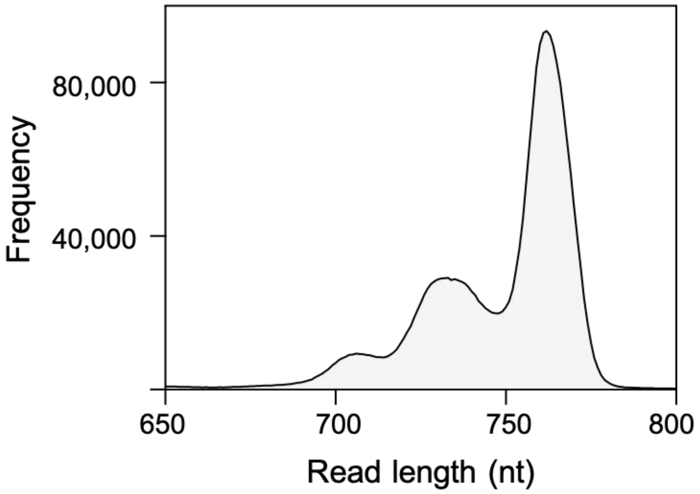


**Figure S4.** The read length profile measured by the nanopore sequencer.

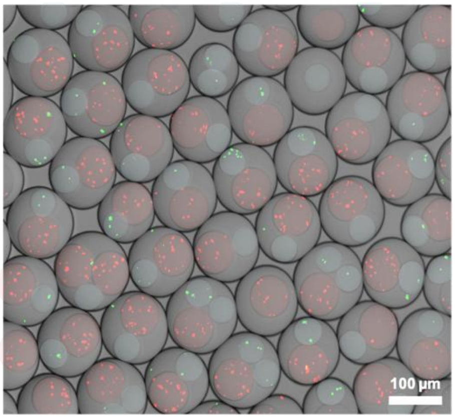


**Figure S5.** Co-encapsulation of hydrogel beads containing bacteria expressing green or red fluorescent proteins within a single droplet.

**Table S1.** Multidimensional comparison between the bead-pairing strategy developed in this work and representative existing bead-loading methods.

| **Technique** | **CelliGO** | **In Drop** | **Microfluidic zipper** | **This work** |
|---|---|---|---|---|
| **Reference** | Gérard A et al. *Nat Biotechnol* **38**:715-721 (2020). | Klein AM et al. *Cell* **161**, 1187-1201 (2015). | Delley CL et al. *Lab Chip* **20**, 2465-2472 (2020). | |
| **1:1 pairing efficiency** | < 1% | < 10% | ~80% | ~90% |
| **Mechanism** | Random encapsulation under double *Possion* distribution | Random encapsulation under *Possion* distribution | Alternated bead close-packing | Deterministic bead encapsulation and synchronized merging |
| **Particle types** | Cell to cell | Cell to bead | Bead to bead | Bead to bead |
| **Co-encapsulation flexibility** | / | / | Fixed at 1:1 | Flexible |
| **Application** | Single-cell activity-based screening | Eukarytotic single-cell transcriptomics | Eukarytotic single-cell DNA profiling | Microbial single-cell 16s rRNA sequencing |
